# The metabolic role of vitamin D in children’s neurodevelopment: a network study

**DOI:** 10.1101/2023.06.23.546277

**Authors:** Margherita De Marzio, Jessica Lasky-Su, Su H. Chu, Nicole Prince, Augusto A. Litonjua, Scott T. Weiss, Rachel S. Kelly, Kimberly R. Glass

## Abstract

Autism spectrum disorder (ASD) is a neurodevelopmental disorder with various proposed environmental risk factors and a rapidly increasing prevalence. Mounting evidence suggests a potential role of vitamin D deficiency in ASD pathogenesis, though the causal mechanisms remain largely unknown. Here we investigate the impact of vitamin D on child neurodevelopment through an integrative network approach that combines metabolomic profiles, clinical traits, and neurodevelopmental data from a pediatric cohort. Our results show that vitamin D deficiency is associated with changes in the metabolic networks of tryptophan, linoleic, and fatty acid metabolism. These changes correlate with distinct ASD-related phenotypes, including delayed communication skills and respiratory dysfunctions. Additionally, our analysis suggests the kynurenine and serotonin sub-pathways may mediate the effect of vitamin D on early childhood communication development. Altogether, our findings provide metabolome-wide insights into the potential of vitamin D as a therapeutic option for ASD and other communication disorders.

## Introduction

Autism spectrum disorder (ASD) is a neurodevelopmental disorder that has increased in prevalence by 176% in the last 20 years^1^. With over 52 million cases globally, it is one of the most rapidly increasing conditions in the world^2, 3^. The etiology of ASD is complex and multifactorial^4–6^. Studies on ASD heritability have identified several genetic risk factors including rare and common genetic variants, chromosomal anomalies, and gene defects^7–9^. Meta-analyses have also identified a series of environmental risk factors, including pre-natal, peri-natal, and post-natal elements^5, 6,10^. Among these factors, vitamin D has been increasingly implicated in ASD pathogenesis^11–13^.

Vitamin D is a fat-soluble vitamin that is produced mainly from skin exposure to UVB radiation^14^. The human body metabolizes vitamin D into a series of steroid-like hormones^15^. These hormones regulate a variety of physiological processes, from bone growth to immune response to brain development and function^15^. Numerous studies have shown associations between ASD and Vitamin D deficiency^11–13^. Children with ASD have lower vitamin D levels in the brain and peripheral blood compared to neurotypical children^16–18^, and children living in areas with low solar UVB doses exhibit three times the ASD prevalence of their peers in sunny areas^19^. During pregnancy, gestational vitamin D deficiency is associated with increased risk of ASD development^20, 21^. Conversely, maternal vitamin D supplementation has been demonstrated to significantly associate with reduced incidence of ASD^22, 23^. Vitamin D supplementation has also shown remarkable therapeutic effects, improving core behavioral and social symptoms of ASD^24–26^.

Multiple modes of action have been proposed to explain the link between vitamin D and ASD^11, 13^. Evidence suggests that vitamin D might be involved in the synthesis of neurotransmitters such as serotonin and dopamine^27, 28^, the modulation of autoimmune responses^29, 30^, and the production of pro-inflammatory cytokines and antioxidants^31, 32^. However, a causal mechanism has yet to be established.

In this study, we employ an innovative and integrative network approach to explore the neurodevelopmental impact of vitamin D on ASD risk. We analyze the metabolomic profiles of 381 children within the Vitamin D Antenatal Asthma Reduction Trial (VDAART)^33^. We investigate how individual metabolic differences translate into differences in vitamin D levels and communication skills, as measured by the Ages and Stages Questionnaire (ASQ)^34, 35^. Early life communication scores derived from this questionnaire are strongly correlated with children’s future ASD risk^36, 37^. The novelty of our work is twofold. Firstly, we present a dataset that uniquely captures information on the metabolomic profile, vitamin D levels, clinical records, and neurobehavioral status of hundreds of children; to our knowledge, no other resources with this combination of information exist. Additionally, we apply network analysis to these data in a novel way. Using LIONESS^38^, a network algorithm developed in our group, we reconstruct each child’s individual metabolic network and integrate these networks with their phenotypic traits.

Collectively, our findings shed light on the functional impact of vitamin D in early childhood communication skills and its association with ASD etiology. By leveraging a metabolome-wide, systems biology approach, this work offers new insights into the potential of vitamin D as a treatment for ASD.

## Results

### The study population

The study population of VDAART has been described previously^33^. Briefly, this study recruited pregnant women at 10 to 18 weeks gestation and randomized them to a daily supplement of 4,000 IU/day of vitamin D3 or placebo. All women also received a daily multivitamin containing 400 IU of vitamin D3. Supplementation took place until delivery. Offspring were evaluated quarterly through questionnaires and yearly through in-person visits.

This study is centered on the clinical and omics profiles of the VDAART children at age three (see Supplementary Table 1). Clinical and phenotypic traits of the children included: demographic information, including race and gender; pathological conditions, including asthma and recurrent wheeze; and developmental predictors collected through the Ages and Stages Questionnaire (ASQ). The ASQ is a parent-completed, developmental screening test evaluating the communication, personal-social, problem-solving, fine motor, and gross motor skills of a child^34, 35^ (see Materials and Methods). For each of these five domains, a child is given a score and categorized as “On Schedule for developing normally” (ASQ=0); “Requires Monitoring” (ASQ=1), or “Needs further evaluation” (ASQ=2). We focused our analysis on the ASQ scores in the communication skills domain (referred to as the ASQ-comm score). This score has been reported to correctly identify 95% of children defined at risk of ASD based on the Modified Checklist for Autism in Toddlers (M-CHAT)^36^. Additionally, we have previously shown that plasma metabolites associated with the ASQ-comm score at age 3 represent accurate predictors of autism diagnosis later in life^37^. For these reasons, we used this score as a proxy for ASD risk in our analyses. All these clinical characteristics were recorded during the annual follow-up visit of children at age three.

During the same year-three visit, children’s blood samples were also collected (see Supplementary Table 1). From these samples, plasma metabolomic profiling was performed by Metabolon Inc. (NC, USA) and serum levels of vitamin D were measured. Serum levels of vitamin D were also measured in mothers before (32 to 38 weeks gestation) and after (1 year) delivery. As such, this dataset provides a unique opportunity to explore the metabolic impact of Vitamin D on postnatal neurodevelopment.

### Reconstructing patient-specific metabolic networks through LIONESS

Prior studies have linked ASD to Vitamin D deficiency during pregnancy and early childhood^12, 13, 39^. The underlying molecular mechanisms remain poorly understood, but a key role for neurotransmitter metabolism has been recently suggested^13, 39^. Building upon these findings, we sought to investigate the metabolic processes associated with the neurodevelopmental actions of vitamin D. To do this, we used LIONESS^38^ (Linear Interpolation to Obtain Network Estimates for Single Samples), a method developed in our group to reverse-engineer sample-specific networks. As opposed to other commonly used network approaches, such as correlation networks, the single-sample networks estimated by LIONESS can model the intrinsic variability across the samples in a data set. This allows us to analyze individual network relationships in connection with individual phenotypes and disease status.

The input to LIONESS is an omics matrix (Fig. 1a). LIONESS estimates the individual network 𝑒^(𝑞)^ of a sample 𝑞 through the following equation^38^ (see also Materials and Methods):

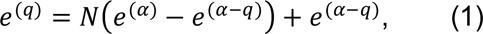

**Fig. 1.**
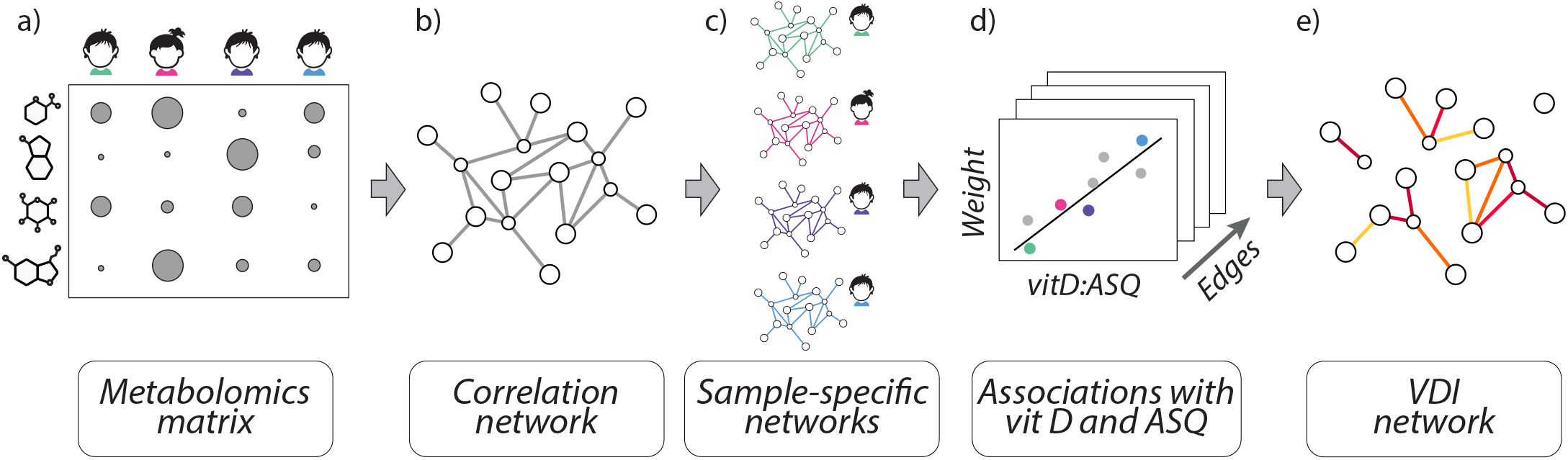
Overall scheme of the analysis. **a)** The metabolomics matrix containing the metabolite levels of 381 VDAART children was used as the input to the LIONESS algorithm. **b)** Pearson correlation coefficients between each pair of metabolites were calculated to build the aggregate correlation network associated with the metabolomics matrix. Only edges with high correlation coefficients are shown for visualization purposes. **c)** LIONESS reconstructed the sample-specific metabolic networks based on Eq. (1). **d)** The LIONESS edge weights for each pair of metabolites were regressed against the children’s phenotypic traits. **e)** We selected edges that are statistically associated with the interaction term between children’s ASQ-comm scores and either maternal or offspring vitamin D levels. These edges constitute the VDI network.

where 𝑁 is the number of samples in the data set (𝛼), and 𝑒^(𝛼)^and 𝑒^(𝛼+𝑞)^are two aggregate networks, built using all samples in the data set (𝑒^(𝛼)^) and built excluding sample 𝑞 (𝑒^(𝛼+𝑞)^) respectively. In our application, we used the metabolomic profiles of the VDAART children at age 3 to build the omics matrix (Fig. 1a). Metabolite levels were filtered for low counts, normalized, corrected for batch effects, and adjusted for covariates (see Materials and Methods). The resulting metabolomic matrix contained the levels of 833 metabolites for 381 samples. We computed the Pearson correlation coefficient for each pair of metabolites to build the two aggregate networks, 𝑒^(𝛼)^ and 𝑒^(𝛼+𝑞)^ (Fig. 1b). Using Eq.(1), we then iteratively applied

LIONESS to reconstruct the metabolic network 𝑒^(𝑞)^ of each VDAART child (Fig. 1c). The edge 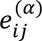 in the individual network 𝑒^(𝑞)^ is a weighted edge connecting two metabolites 𝑖 and 𝑗. The edge’s weight represents how much sample 𝑞 contributes to the co-expression between the metabolites 𝑖 and 𝑗. The more positive (negative) 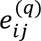, the more the pair (𝑖, 𝑗) has coordinated (anti-coordinated) metabolic expression in the sample 𝑞. In total, LIONESS generated 381 individual networks, each composed of 833 nodes and 346,528 edges (Supplementary Table 2). These networks allowed us to investigate the relationship between the children’s metabolic profiles and their phenotypes.

### Tryptophan, linoleic, and fatty acid metabolism are associated with the neurodevelopmental role of vitamin D in early childhood

Starting from the LIONESS networks, we examined the metabolic interactions associated with the role of vitamin D in children’s neurodevelopment. To do so, we ran a linear regression analysis on the children’s metabolic networks against their phenotypic traits (Fig. 1d). For each metabolite pair, the weight of the associated LIONESS edge across the individual networks was regressed against multiple clinical variables (Fig. 1d, see Materials and Methods). We selected metabolite edges that were significantly associated with 1) the interaction term between children’s ASQ-comm score and their Vitamin D levels, and/or 2) the interaction term between children’s ASQ-comm score and their mother’s Vitamin D levels at 32 to 38 gestation weeks (p-value < 0.01). We reasoned that these edges represent potential metabolic reactions affected by vitamin D in relation to communication skills at age 3. Therefore, they might highlight biochemical processes mediating the interaction between Vitamin D and ASD. We refer to the subnetwork composed of these significant edges as the Vitamin D Interaction (VDI) network (Fig. 1e, Supplementary Table 3).

To determine the metabolic pathways underlying the VDI network, we performed a pre-ranked Metabolic Set Enrichment Analysis (MSEA)^40, 41^ using the Kyoto Encyclopedia of Genes and Genome (KEGG)^42^ (Fig. 2a). We ranked all the metabolites based on their number of connections, or “degree”, in the VDI network and ran MSEA based on this pre-ranked list (see Materials and Methods). In this way, the MSEA was built upon the topology of the VDI network: enriched pathways represent molecular processes where the annotated metabolites act predominantly as hubs in the VDI network (nodes with a large number of edges). Among the most enriched metabolic pathways (Supplementary Table 4), we identified Tryptophan metabolism (hsa00380, FDR=0.08), Linoleic Acid Metabolism (hsa00591, FDR=0.002), and Biosynthesis of Unsaturated Fatty Acids (hsa01040, FDR = 10^-6^).

**Fig. 2.**
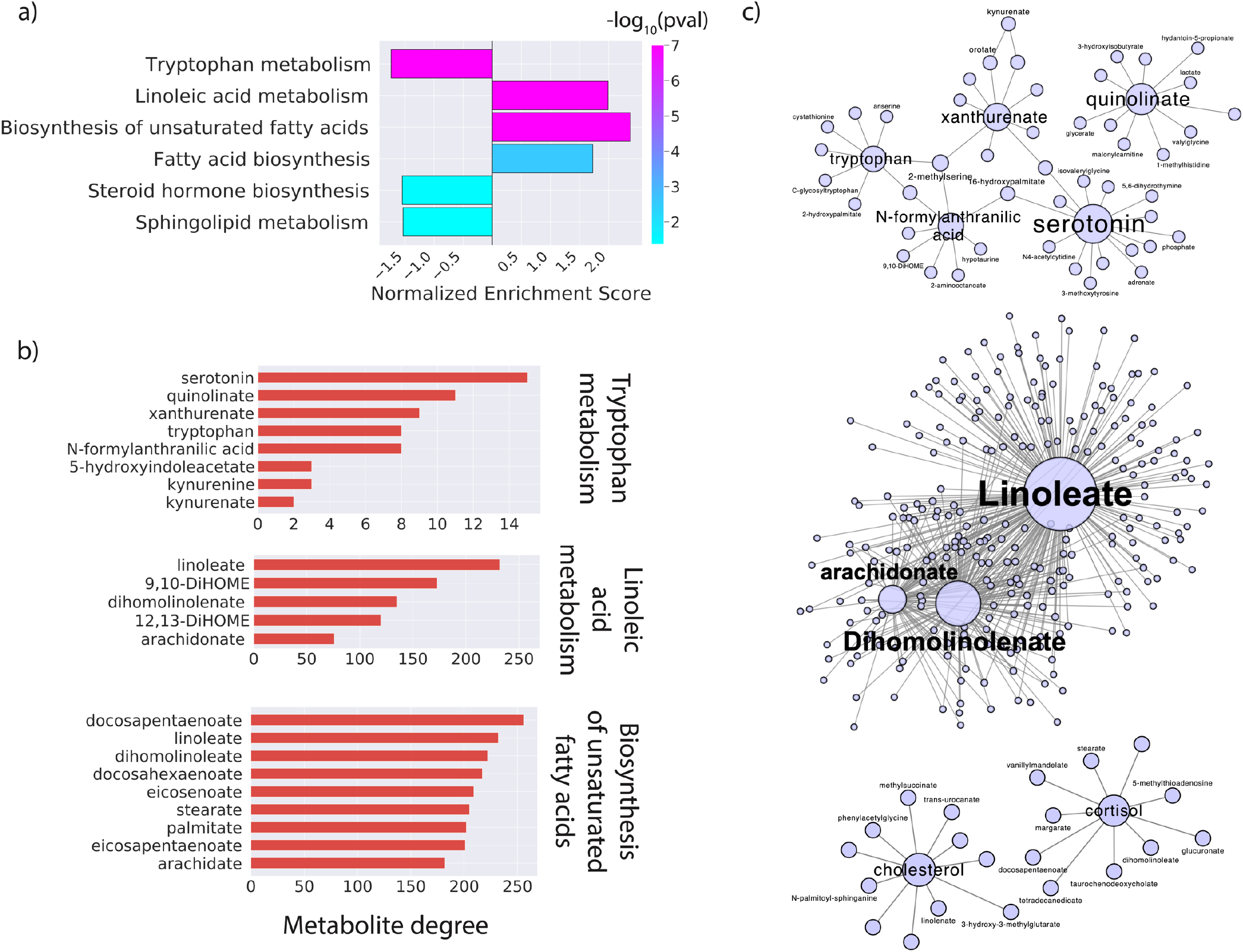
Enrichment analysis of the VDI network. **a)** Normalized Enrichment Score of KEGG pathways enriched in the VDI network based on pre-ranked MSEA. Pathways are ranked based on their p-values. **b)** VDI network degree of the MSEA leading metabolites of the top three enriched KEGG metabolic pathways, which include Tryptophan metabolism, Linoleic acid metabolism, and Biosynthesis of unsaturated fatty acids. **c)** First neighbors’ network of the leading metabolites for 1) Tryptophan metabolism (top), 2) Linoleic acid metabolism (center), and 3) Steroid hormone biosynthesis (bottom). Node sizes are proportional to the node’s degree. Only edges included in the VDI network are shown. For visualization purposes, we highlighted nodes that were mentioned in the main text.

Within the Tryptophan pathway, MSEA leading edge analysis^43^ highlighted serotonin, quinolinate, xanthurenate, tryptophan, and kynurenine as leading metabolites (Fig. 2b and 2c). Interestingly, altered regulation of these neuroactive metabolites has been previously linked to ASD^44–46^.

The Linoleic Acid Metabolism pathway also emerged as highly enriched in our VDI network. MSEA leading metabolites in this pathway included linoleate, dihomolinolenate, and arachidonate (Fig. 2b). These metabolites represented large hubs in the VDI network, with over 70 metabolite edges significantly associated with the interaction between vitamin D and the ASQ-comm score (Fig. 2c). Multiple studies have reported deficiency of these metabolites in individuals with ASD^47–49^. This suggested that vitamin D may impact lipid metabolism, which is known to affect brain growth and cognitive development^47, 48^.

The VDI network was enriched in pathways related to fatty acid production. This result supported the growing evidence of ASD as a disorder of fatty acid metabolism^49–51^. Leading metabolites annotated to the Biosynthesis of Unsaturated Fatty Acids included docosahexaenoate, stearate, and palmitate (Fig. 2b), whose levels have been shown to be reduced in individuals with ASD^49^. Docosahexaenoate was a major hub in our VDI network (Fig. 2b). It’s worth noting that vitamin D and docosahexaenoic acid have been proposed to act synergistically in modulating serotonin synthesis, which may help ASD symptoms^52^. In conjunction with aberrant metabolism of unsaturated fatty acids, a deficit in cholesterol biosynthesis may be involved in the pathogenesis of ASD^51^. Cholesterol was a leading metabolite in the enrichment of the Steroid Hormone Biosynthesis pathway (Fig. 2c). It follows that vitamin D deficiency might affect neurodevelopment by disrupting cholesterol homeostasis.

### Metabolic interactions of Vitamin D with children’s neurodevelopment correlate with different disease outcomes

One of the clinical challenges of ASD’s diagnosis is the heterogeneity of its symptomatology^53, 54^. Our network analysis suggested that Vitamin D may affect ASD development through multiple metabolic pathways. Therefore, we interrogated whether differences within these pathways underlie epidemiological differences among the VDAART children. To answer this question, we performed hierarchical clustering of the VDAART children based on the LIONESS weights of the VDI edges in their individual networks (see Materials and Methods). Our analysis identified 5 main clusters. Notably, each cluster exhibited different clinical, physiological, and metabolic characteristics (Supplementary Table 5).

Children’s vitamin D levels were overall higher in cluster 3 and 5, with cluster 5 having significantly higher vitamin D levels at age 3 compared to all other clusters (Student’s T-test, P = 0.039; see Material and Methods for details). In contrast, children in cluster 4 exhibited significantly lower vitamin D (Student’s T-test, P = 0.062; Fig. 3a). Interestingly, maternal vitamin D levels collected during and after pregnancy exhibited a behavior similar to the offspring (Fig. 3a). On average, clusters 3 and 5 showed higher maternal vitamin D throughout all the measured endpoints. Mothers of the children in cluster 3 had significantly higher vitamin D both during late pregnancy (32-38 gestation weeks, Student’s T-test, P = 0.09) and at one year after delivery (Student’s T-test, P = 0.1), while children in cluster 5 had significantly higher cord blood vitamin D (Student’s T-test, P = 0.085). This result suggested that imbalances in maternal vitamin D may lead to imbalances in vitamin D during the offspring’s early childhood. Notably, this interdependence was not evident when looking purely at the correlation between the mothers’ and children’s vitamin D measurements (Supplementary Figure 1).

**Fig. 3.**
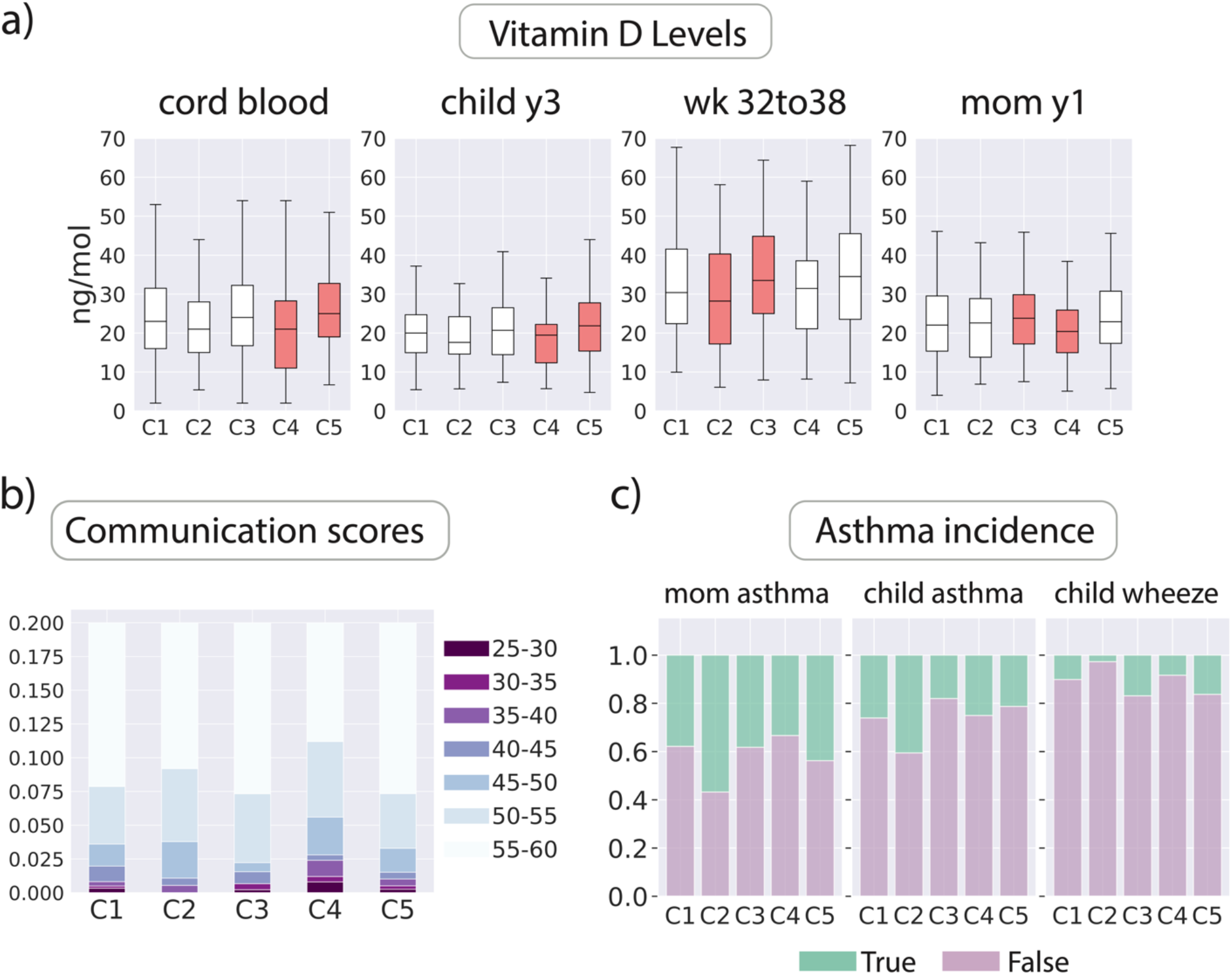
Children’s phenotypic traits in the clusters extracted from the VDI network. **a)** Vitamin D levels (ng/mol) of children in each cluster at year 3, their mothers at 32 to 38 gestation weeks and one year after delivery, and in the cord blood. Colored boxes represent clusters with lower or higher Vitamin D levels compared to the individuals in all other clusters (p-value<0.15, see Materials and Methods) **b)** Stacked histogram of the communication scores for children in each cluster. Cluster 4 (referred as the low-communication cluster) exhibited the lowest distribution of communication scores. **c)** Stacked histogram of asthma incidence in children and mothers within each cluster. Cluster 2 (referred as the asthma cluster) exhibited the highest proportion of children and mothers with asthma.

We next analyzed the VDAART communication scores obtained from the ASQ questions related to child’s communication skills (see Materials and Methods). These scores reflect a child’s proficiency in different aspects of communication; the higher the score, the higher the child’s ability to perform communication tasks. The distribution of communication scores across clusters revealed a pattern opposite to that of the vitamin D levels (Fig. 3b). Children in clusters 3 and 5 exhibited higher communication scores, with cluster 3 reaching statistical significance (Mann-Whitney U test, P = 0.04). Children in cluster 4 showed significantly lower scores (Mann-Whitney U test, P = 0.015). As such, cluster 4 represent children with delayed communication development compared to the rest of the population. We refer to this cluster as the low-communication cluster. The lower levels of vitamin D in this cluster supported a neuroprotective role of vitamin D in early childhood communication development.

The VDAART cohort was initially established to examine the impact of vitamin D on asthma and allergies^33^, and therefore includes information regarding children’s current and family history of asthma and respiratory symptoms. Considering the elevated likelihood of children with ASD to have asthma^55, 56^, we investigated potential connections between our VDI network and the incidence of asthma. Fig. 3c displays the VDAART children’s symptomatology together with their parental history. Comparison between clusters showed a significantly higher incidence of asthma and maternal asthma in cluster 2 (Chi-Squared Test, P= 0.03, 0.05 for child’s and maternal asthma respectively). Together with the low-communication cluster, this cluster also exhibited the lowest maternal vitamin D levels during late pregnancy (Student’s T-test, P = 0.148). At age three, 70% of children in cluster 2 had Vitamin D concentrations below the deficiency threshold of 20 ng/ml^57^. As such, cluster 2 (referred to as the asthma cluster) reflect a different phenotypic outcome associated with vitamin D deficiency. This result can be interpreted within the context of the immunomodulatory and anti-inflammatory effects of vitamin D. Vitamin D might alter the metabolic regulation of the immune response, leading to the respiratory abnormalities often encountered in individuals with ASD. In support of this, children in cluster 3 had lower incidence of asthma (Chi-Squared Test, P=0.14).

We further examined if the different disease phenotypes of the low-communication and asthma clusters coincided with different metabolic endotypes. To do so, we compared the clusters’ individual metabolic networks and identified their marker metabolic edges (see Materials and Methods). For each cluster, a marker edge is defined as an edge whose LIONESS weights significantly differ in that cluster compared to the rest of the population. In agreement with our enrichment analysis, marker edges of the low-communication cluster involved interactions with serotonin and its breakdown product 5-hydroxyindoleacetate (5HIAA) (Fig. 4a). Top significant edges connected 5HIAA to ester derivatives of L-carnitine, including 3-Hydroxybutyrylcarnitine, acetyl-L-carnitine, and palmitoleoylcarnitine. This hinted at a role of vitamin D in the crosstalk between serotonin synthesis and the carnitine shuttle system^58^. Of note, ASD patients often present with carnitine deficiency^59, 60^. Marker edges of the low-communication cluster also involved interactions between 5HIAA and members of the linoleate metabolism pathway, suggesting perturbations in fatty acid production. Additionally, this cluster exhibited altered connection with the amino acid proline (Fig. 4a). In support of this result, abnormal proline levels have been associated with the catechol-O-methyltransferase genotype, a gene variant affecting the brain dopaminergic system in individuals with ASD^61^. Finally, marker edges involved interactions with palmitic and stearic acids, which are both ASD biomarkers^48, 49^.

**Fig. 4.**
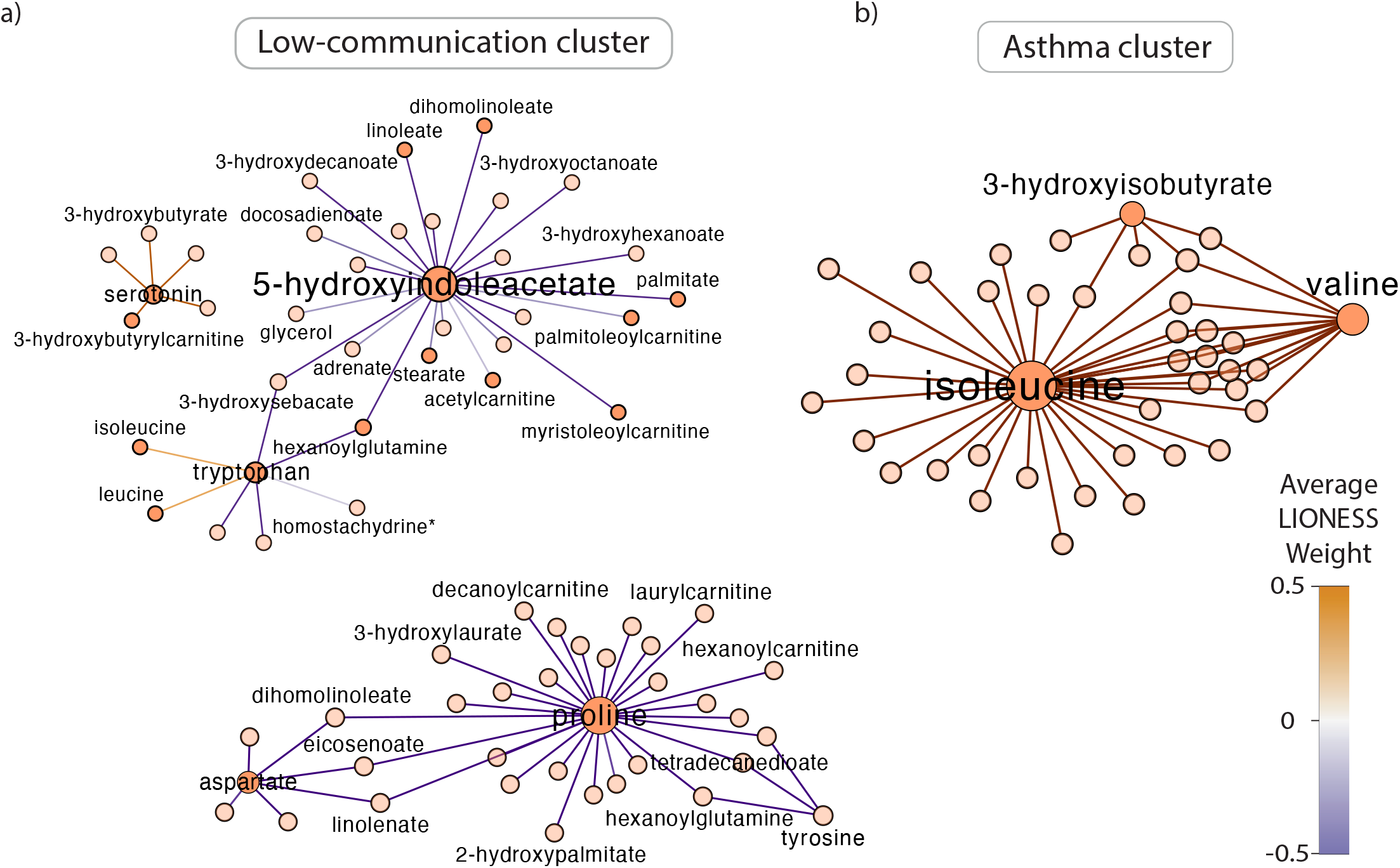
Marker metabolic edges identified for the low-communication and asthma clusters. Top marker metabolic edges identified for the **a)** low-communication and **b)** asthma clusters. Node sizes are proportional to the node’s degree in the marker edges’ subnetwork. Edges are colored based on the average LIONESS weight across the networks in the cluster. For visualization purposes, we highlighted nodes and edges that were mentioned in the main text.

Analysis of marker edges of the asthma cluster revealed a different pattern. Several edges involved the amino-acids isoleucine and valine (Fig. 4b). This result was noteworthy; previous reports have listed these metabolites as differentially expressed in the plasma metabolome of asthma and asthma-COPD overlap patients^62, 63^. The metabolic networks of the asthma cluster also showed perturbations in the palmitic and stearic acid metabolism pathways.

These findings indicated that vitamin D affects children’s neurodevelopment through multiple mechanisms of action, leading to the development of distinct metabolic endotypes and phenotypes.

### The VDI network highlights a role of the kynurenine and serotonin sub-pathways in the interaction between vitamin D and ASD

Our MSEA analysis revealed tryptophan metabolism as one of the most enriched pathways within the VDI network (Fig. 2a and Supplementary Table 4). Our previous studies^37^ have also identified strong associations between tryptophan metabolites and the ASQ-comm score in both the plasma and stool metabolome of VDAART children. Given this evidence, we investigated the biochemical reactions underlying these associations. We selected metabolic edges that were 1) statistically associated with the interaction between children’s and maternal vitamin D and ASQ-comm score (p-value<0.05) and 2) connecting pairs of metabolites that are both annotated to the KEGG tryptophan metabolism pathway. We identified only one edge connecting 5HIAA and L-kynurenine. This edge was also significantly associated with children’s vitamin D levels in our regression model (p-value = 6.4 × 10^-3^). By definition, the metabolites of this edges exhibited co-expression changes in association with the interaction between vitamin D and the ASQ-comm score. To identify the molecular mechanisms subtending these changes, we mapped 5HIAA and L-kynurenine onto the KEGG tryptophan reaction network (see Materials and Methods). The nodes of this network represent all chemical compounds listed in the KEGG tryptophan metabolism pathway. The edges represent chemical reactions between two metabolites, a substrate and a product.

To gain insights on the cascade of biochemical reactions affecting the relative proportion of 5HIAA and L-kynurenine, we computed all the shortest paths connecting these two metabolites within the tryptophan reaction network. We identified one path (Fig. 5), involving two main subprocesses: 1) the biosynthesis of serotonin from L-tryptophan and 2) the degradation of L-tryptophan via the kynurenine pathway. Biochemical reactions mediating serotonin biosynthesis included hydroxylation of L-tryptophan into L-5-hydroxytryptophan (5-HTP) (KEGG reaction R01814), subsequent decarboxylation of 5-HTP to produce serotonin (KEGG R02701), serotonin degradation into 5-hydroxyindoleacetaldehyde(5-HIAL) (KEGG R02908), and the final oxidation of 5-HIAL to 5HIAA (KEGG R04903) (Fig. 5). Our shortest path also contained metabolic reactions involved in the kynurenine pathway, including the oxidation of L-tryptophan to N-formylkynurenine (NFK) (KEGG R00678) and subsequent hydrolysis of NFK to L-kynurenine (KEGG R01959). For each of these biochemical reactions, we analyzed their substrate-to-product expression ratio in association with the children’s ASQ-comm score (see Materials and Methods). The ratio between serotonin and 5HIAA levels increased significantly with increasing ASQ-comm score (p-value= 0.038), suggesting dysregulated activation of serotonin biosynthesis in children with impaired communication skills. This result was consistent with previous observations of blood hyper-serotoninemia in children with ASD^27, 64^.

**Fig. 5.**
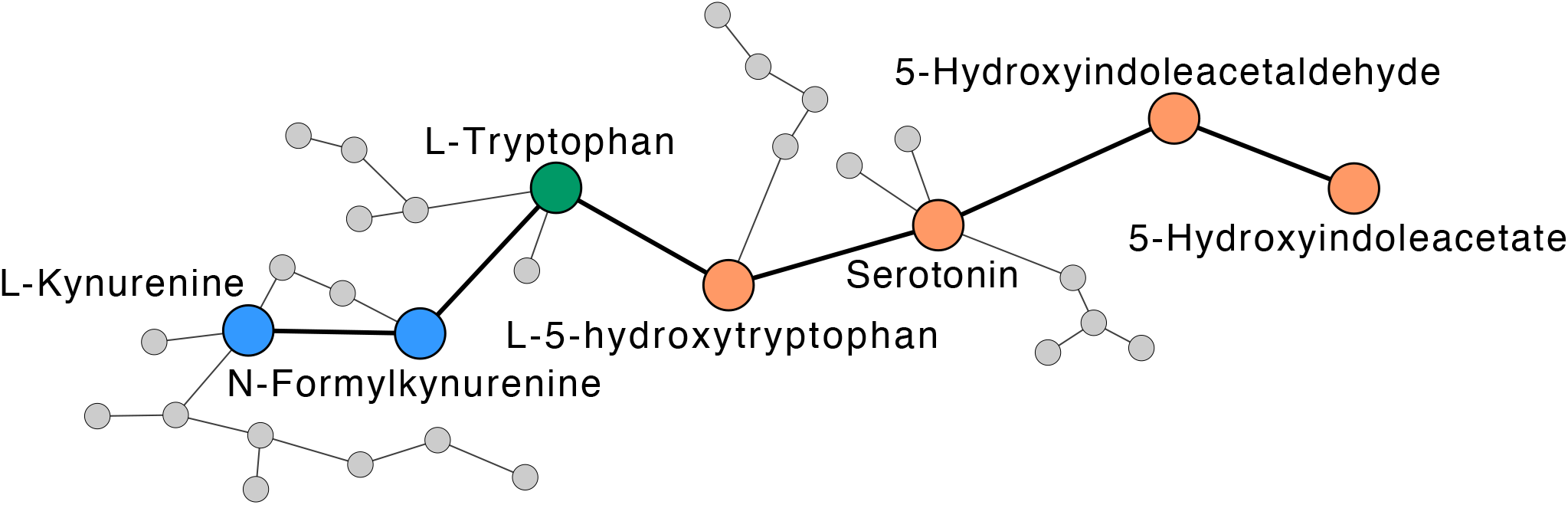
Shortest path connecting 5HIAA and L-kynurenine in the KEGG reaction network. The edge connecting 5HIAA and L-kynurenine was statistically associated with the interaction term between vitamin D and the ASQ communication score. One shortest path connected these two metabolites in the KEGG reaction network of tryptophan metabolism. This path involved two main subprocesses: 1) the biosynthesis of serotonin from L-tryptophan (orange nodes) and 2) the degradation of L-tryptophan via the kynurenine pathway (blue nodes). The nodes in this network represent chemical compounds listed in the KEGG tryptophan metabolism. The edges represent the chemical reactions between a substrate and a product. Only the largest connected component is shown.

## Discussion

This work investigates the impact of vitamin D on the human metabolic network in association with children’s communication development. Language and communication deficits represent a characterizing feature of ASD^4–6^, and the assessment of communication impairments has been proposed as an effective screening for ASD risk^36, 37, 65^. By using early childhood communication skills as a proxy for ASD, our analysis sheds light on the potential neuroprotective effect of vitamin D against ASD.

We integrated the metabolomic profiles, clinical traits, and communication screening data of hundreds of children through a unique network framework. This is the first instance where our LIONESS method has been applied in the context of paired metabolomics and phenotypic data, illustrating the novelty of our approach. Our results suggest that the neurodevelopmental action of vitamin D operates through several metabolic pathways, altering interactions within the tryptophan metabolism, lipid metabolism, and unsaturated fatty acid metabolism pathways. Closer analysis of the tryptophan pathway highlights a key role of the kynurenine and serotonin sub-pathways. These findings provide new mechanistic insights in support of known statistical associations between vitamin D and ASD.

A substantial body of literature has linked ASD to disrupted levels of neuroactive metabolites^27, 64, 66–69^: blood hyper-serotoninemia is one of the most consistent biomarkers of ASD^27, 64^; overproduction of quinolinic, xanthurenic, and kynurenic acid has been observed in individuals with ASD and has been linked to enhanced kynurenine and glutamatergic activity^68, 69^; finally, vitamin D has been proposed to differentially regulate the transcription of the serotonin-synthesizing genes *TPH1* and *TPH2*^27^. In support of these findings, our network analysis suggests that vitamin D might play a role by altering metabolic interactions between serotonin and other members of the tryptophan metabolism pathway, including xanthurenate, L-tryptophan, N-formylanthranilic acid, and quinolinate. Additionally, ASD has been associated with fatty acid deficiencies^49–51, 70^. Here we show that differences in vitamin D levels correspond to perturbations in fatty acid metabolism, including the metabolic interactions between docosahexaenoate, stearate, and palmitate. Interestingly, these metabolites are major constituents of membrane phospholipids. This supports recent mechanistic theories about dysregulation of phospholipid metabolism in patients affected by neurodevelopmental disorders, including ASD and attention-deficit/hyperactivity disorder^71, 72^. Finally, ASD manifestations are often accompanied by other comorbid pathologies^55, 56^. Our findings suggest that vitamin D deficiency can impact different components of children’s metabolic networks. These components are associated with different disease outcomes, including impaired communications skills and asthma.

There are at least three postulated mechanisms by which vitamin D might be related to ASD. Firstly, vitamin D might be critically involved in serotonin metabolism and vitamin D deficiency could result in reduced brain serotonin levels, as observed in ASD patients^11, 13, 27, 66^. Secondly, the anti-inflammatory properties of vitamin D3 may serve as a neuroprotective mechanism against ASD by directly reducing neurotoxin levels in the brain^11, 13, 73^. Thirdly, vitamin D deficiency combined with excess androgens may contribute to ASD development through potential interactions with sex hormones^11, 74–77^. These theories are not mutually exclusive. Our data speak only to the first hypothesis and not to the other two possibilities.

Our work has some limitations. We recognize that the ASQ communication score does not represent a clinical diagnosis of ASD. Indeed, there are intrinsic biases when estimating children’s developmental status from indirect caregiver reports such as the ASQ^36, 78^. However, our dataset represents one of the first instances where paired information on the metabolic profile, vitamin D levels, and neurobehavioral status of hundreds of children has been captured. As such, this study is only the first step towards determining the role of vitamin D on neurodevelopment in early childhood. Further clinical and translational assessments will be crucial for establishing a definite link between vitamin D and ASD.

Our findings provide a comprehensive metabolome-wide, systems-biology perspective on the molecular effects of vitamin D on children’s neurodevelopment. Nevertheless, questions remain regarding the reproducibility of our biological findings. Future experiments and cross-validation analyses are needed to address these questions, laying the groundwork for the implementation of clinical trials investigating the potential of vitamin D in ASD treatment.

## Materials and Methods

### Calculation of the ASQ-comm score

Primary caregivers of VDAART children submitted the Ages and Stages Questionnaire^35^ (3rd Edition, https://agesandstages.com/; Paul H. Brookes Publishing Co., Inc.) during the children’s annual three year visit. In the ASQ, caregivers were asked a series of questions about the ability of their children to perform a particular task. Answers to each question were scored with 10 points if the answer was “Yes”, 5 points if the answer was “Sometimes”, and 0 points if the answer was “Not yet”. Questions pertained to five main developmental domains: gross motor skills, fine motor skills, problem solving ability, personal/social skills, and communication. Six questions per domain were assigned, resulting in a domain specific score. In this work, we focused on the scores from the communication skills domain. We referred to this score as the “communication score”.

The communication score was compared to the expected mean score of a reference distribution within age-groups. Scores were then categorized as follows: “On Schedule for developing normally”; “Requires Monitoring” (1–2 standard deviations below the mean); and “Needs further evaluation” (>2 standard deviations below the mean). We referred to this categorization as the ASQ-comm score.

### Metabolomic profiling and preprocessing

We analyzed previously generated and preprocessed metabolomic data^37^. Briefly, blood samples were obtained from participating children in VDAART at age three years. Metabolites were analyzed as measured LC-MS peak areas and identified by their mass-to-charge ratio, retention time, and through a comparison to library entries of purified known standards. The blood samples were processed in two batches sent six months apart (batch one n = 245; batch two n = 688) then merged and scaled together based on equivalence of the control groups. If a metabolite was missing in 50% or more of the samples from either dataset, it was excluded from further analysis. All remaining missing values were imputed with half the minimum peak intensity for that metabolite across the whole population. Data were pareto scaled to account for the differences in the scales of measurements across the metabolome. Metabolites were log-transformed to create approximately Gaussian distributions and to stabilize variance. See our previous paper^37^ for extensive description of data extraction and processing.

### LIONESS

Correlation networks have been successful in aggregating and summarizing multidimensional biological data^79, 80^, but they do not account for the intrinsic variability across the samples of a population. To fill this gap, our group developed LIONESS^38^, a method to reverse-engineer sample-specific networks starting from the aggregate network of an entire population.

The founding principle of LIONESS is that an aggregate network, summarizing information from a set (𝛼) of 𝑁 samples, can be conceptualized as the average of individual component networks, representing the contributions from each member within the input sample set. This principle can be formalized mathematically through the following equation:

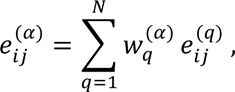

where 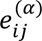 and 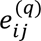 represent a weighted edge between two nodes 𝑖 and 𝑗 in the set-wide aggregate network and in the individual network of sample 𝑞, respectively. 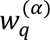 represents the contributing weight of sample 𝑞 to the set (𝛼). If all samples have equal weight, 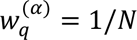. Starting from the above equation, the individual network 𝑒^(𝑞)^of sample 𝑞 can be reconstructed as (see original paper^38^ for derivations):

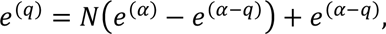

where 𝑒^(𝛼)^is the aggregate network estimated using all 𝑁 samples in the set (𝛼) and 𝑒^(𝛼+𝑞)^is the aggregate network estimated using all samples except for sample *q*. In these aggregate networks, nodes may represent individual omics entities (e.g. metabolites) and edges may represent relationships summarizing biological information from the population-based omics matrix (e.g. the correlation between two metabolites). Intuitively, this equation expresses an individual network as the sum of two contributions: one that accounts for biological patterns that are specific to the individual 𝑞, and a second that accounts for biological patterns that are shared across all the samples in the population.

In our application, we employed Pearson correlation to build the two aggregate metabolic networks 𝑒^(𝛼)^ and 𝑒^(𝛼+𝑞)^for the VDAART study sample (𝛼). In these correlation networks, each metabolite pair (𝑖, 𝑗) is connected through a weighted edge. The edge’s weight represents the co-expression coefficient between 𝑖 and 𝑗 calculated across the VDAART study sample before (𝑒^(𝛼)^) and after (𝑒^(𝛼+𝑞)^) removing individual 𝑞. Using Eq.(1), we reconstructed the individual weighted network of each VDAART child.

### Linear regression model to generate the VDI network

LIONESS generated 381 fully connected, individual, weighted networks. For each edge in those networks, we regressed the distribution of edge weights across the VDAART children against their phenotypic traits and family history. We used the following multivariable regression model:

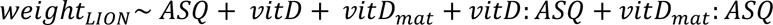

The terms 𝐴𝑆𝑄 and 𝑣𝑖𝑡𝐷 represent children’s ASQ-comm score and vitamin D levels at age three years, respectively; 𝑣𝑖𝑡𝐷_𝑚𝑎𝑡_represents maternal Vitamin D levels during late pregnancy (within 32-38 weeks); and 𝑣𝑖𝑡𝐷: 𝐴𝑆𝑄 and 𝑣𝑖𝑡𝐷_𝑚𝑎𝑡_: 𝐴𝑆𝑄 represent interaction terms between the ASQ-comm score and children’s and maternal vitamin D levels, respectively. This model allows us to identify the metabolic edges associated with the interaction between maternal and early childhood vitamin D levels and child’s communication development. The model was adjusted for the children’s race and sex, maternal education level and marital status, treatment group, and clinical site of plasma collection (see Supplementary Table 1). Linear regression was performed using the function *lmfit* in the R package LIMMA^81^. P-value statistics were obtained from the LIMMA function *eBayes*. This analysis provided a list of linear regression coefficients and p-values for each edge (metabolite pairs).

### Pre-ranked MSEA Enrichment analysis

MSEA enrichment analysis was performed using the python package GSEApy^82^. First, we ranked all metabolites in the VDI network based on their degree. If two nodes had the same degree, the one having the highest “Interaction Coefficient” (IC) was ranked higher. We defined the IC of a node as the mean absolute interaction coefficient 𝑚𝑒𝑎𝑛(𝑣𝑖𝑡𝐷: 𝐴𝑆𝑄, 𝑣𝑖𝑡𝐷_𝑚𝑎𝑡_: 𝐴𝑆𝑄) averaged across all the VDI edges connected to the node, as calculated from the linear regression analysis. Nodes with high IC exhibit connections highly associated with the interaction terms between maternal and children’s vitamin D and the ASQ-comm score. As such, they likely represent key regulators of these interactions. Nodes that were not included in the VDI network were assigned degree zero. To break ties between zero-degree nodes, we ranked them based on their IC. We downloaded all the human metabolic pathways within the KEGG^42^ database using the functions *process_kegg* and *process_gmt* of the python package ssPA^40^. Based on these KEGG pathways, we performed pre-ranked Metabolic Set Enrichment Analysis (MSEA) on our pre-ranked list of VDI metabolites using the function *prerank* of GSEApy. P-values were adjusted using the Benjamini-Hochberg (BH)-FDR correction, and an FDR < 0.15 was used to identify significantly enriched pathways.

### Hierarchical clustering analysis of the individual metabolic networks

To identify similarities in the metabolic networks of the VDAART children, we selected the subgraph of VDI edges in each LIONESS network. We then z-score normalized the weights of each edge across the individual VDI networks and performed hierarchical clustering on the z-scores using Spearman correlation as similarity distance and the complete-linkage metrics. Based on the hierarchical structure of the cluster dendrogram, the optimal number of clusters was obtained by cutting the tree such that all the descendent links in each cluster are shorter than a cut-off distance of 1.15. We obtained five clusters (see Supplementary Figure 2). We compared the phenotypic and molecular characteristics of these clusters.

In each cluster, we analyzed 1) children’s communication scores and vitamin D levels at age three, 2) maternal vitamin D levels during pregnancy (within 32-38 weeks), in the cord blood, and 1 year after delivery, and 3) children’s history of asthma/wheeze, and 4) maternal asthma. For each of these variables, we compared its distribution in each cluster versus all other clusters. P-value statistics were obtained using the Mann-Whitney u test for the communication score, T-test for the other numerical variables, and chi-squared for binary variables. A cluster was considered significantly different from the others in a given variable if its p-value was less than 0.1. For reference, in the main text we reported p-values trends lower than 0.15.

Additionally, we identified marker metabolic edges for each cluster. For each edge connecting a pair of metabolites, we used a T-test to compare the distribution of that edge’s LIONESS weight in the cluster versus the rest of the population. An edge was considered a marker edge in the cluster if its p-value was less than 0.1.

### KEGG tryptophan reaction network

To build the tryptophan reaction network, we used the Biopython^83^ package Bio.Kegg. We downloaded the KEGG Markup Language^42^ (KGML) associated with the tryptophan metabolism pathway (hsa00380). We then selected the chemical network from the KGML graph object. In this network, nodes represent chemical compounds and edges represent chemical reactions between a substrate and a product (as defined in the KEGG documentation^42^). The resulting reaction network was composed of 41 nodes and 39 edges. We mapped all the metabolites of our VDI networks on to the KEGG reaction network and computed the shortest paths between pairs of metabolites connected by a VDI edge. We identified one shortest path connecting L-kynurenine with 5HIAA. For each metabolite pair (𝐴, 𝐵) in this path, we investigated if its expression ratio was statistically associated with the children’s ASQ-comm scores. Specifically, we used the linear regression model 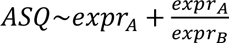 . The model was adjusted for the children’s race and sex, maternal education level and marital status, treatment group, and clinical site of plasma collection.

## Author contributions

M.D.M. analyzed the data. M.D.M., S.T.W., R.S.K., and K.R.G. interpreted the data and wrote the manuscript. All the authors contributed to the writing of the paper, provided critical feedback, and helped shape the research and the analysis of the problem.

## Data and materials availability

All data needed to evaluate the conclusions in the paper are present in the paper and/or the Supplementary Materials. Additional data related to this paper may be requested from the authors.

## Supplementary Material

### Supplementary Figures

**Supp. Fig. 1.**
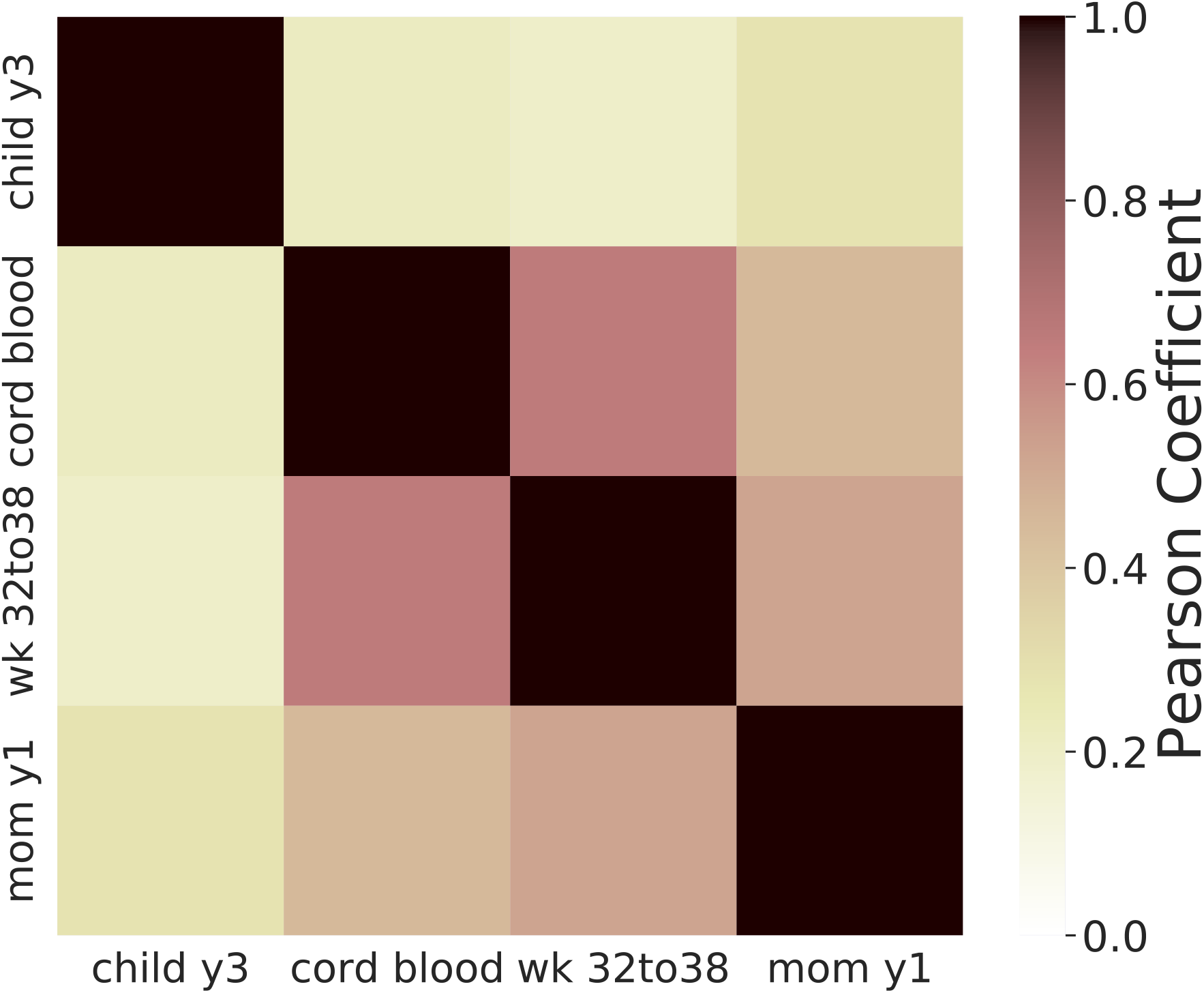
Correlation between maternal and children’s vitamin D levels. Heatmap of the Pearson correlation coefficients between children’s vitamin D levels at age three and their mothers during and after pregnancy. Without segregating children in separate clusters, no evident correlation is observed between children’s and maternal vitamin D levels across the entire VDAART population.

**Supp. Fig. 2.**
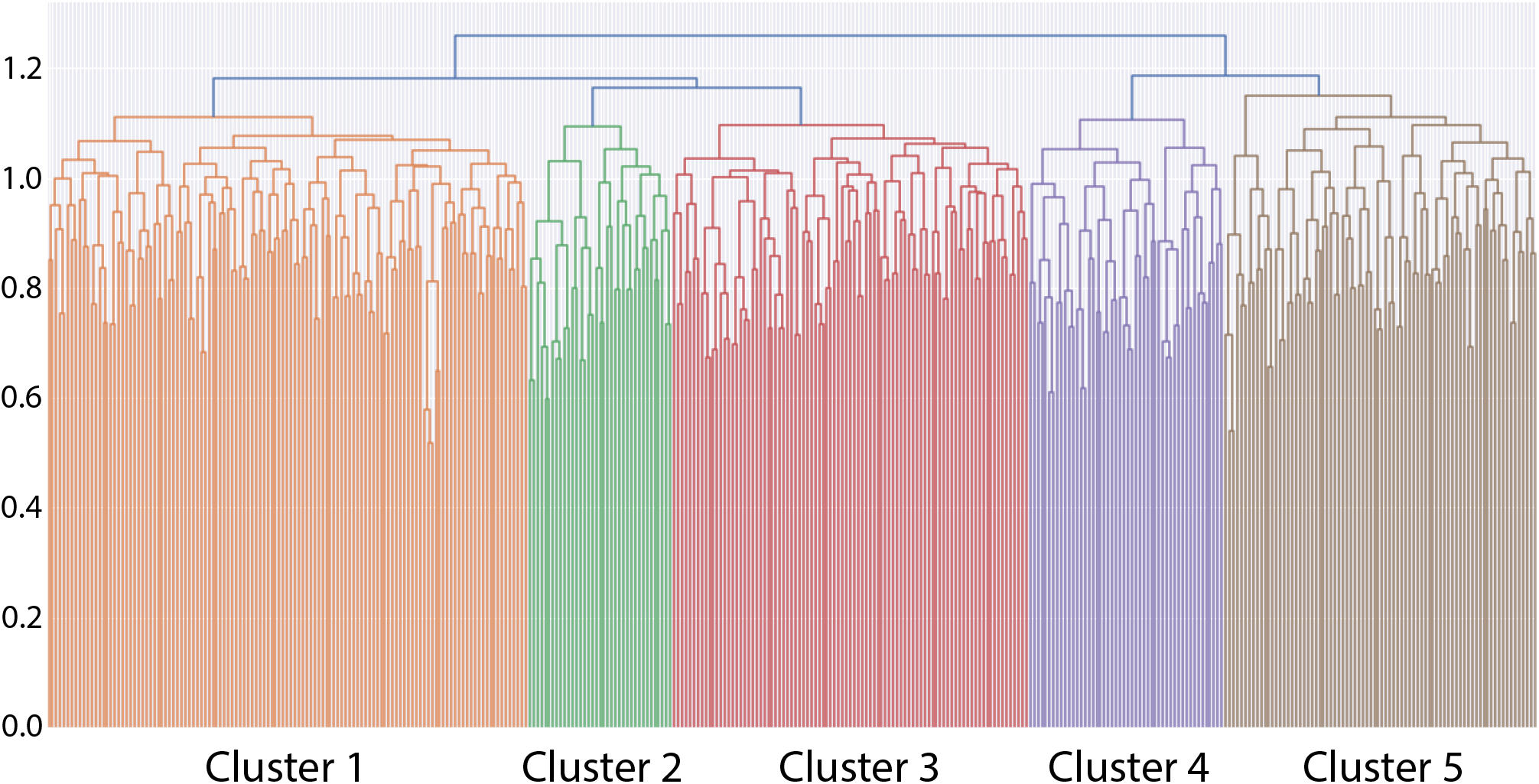
Dendrogram of the hierarchical clustering of VDI edges in the LIONESS networks. The height of each link in the dendrogram (y-axis) is the distance metric, calculated as 1 – the Spearman correlation. The dendrogram was constructed using complete-linkage. Cutting the dendrogram at the height of 1.15 results in 5 clusters.

### Supplementary Tables

**Supplementary Table 1.** (Top) Baseline characteristics of the 381 children from the Vitamin D Antenatal Asthma Reduction Trial (VDAART) studied in this manuscript. (Bottom) Summary of plasma metabolomic profiling, serum vitamin D levels, and Ages and Stages Questionnaire (ASQ) scores for these children and their mothers.

**Supplementary Table 2.** The weights of the 346,528 metabolic edges predicted by LIONESS for each individual network of the 318 VDAART children.

**Supplementary Table 3.** A list of the VDI edges together with their associated coefficients and the p-values resulting from the linear regression model.

**Supplementary Table 4.** A list of the KEGG metabolic pathways that are enriched in the VDI network based on our MSEA pre-ranked enrichment analysis. For each pathway, we included its KEGG ID, Enrichment Score (es), Normalized Enrichment Score (nes), nominal p-value (pval), FDR-adjusted p-value (fdr), number of metabolites in the pathway (set_size), number of metabolites in the pathway matched to the data (matched_size), metabolites in the pathway matched to the data (Pathway_metabolites), and leading edge metabolites (leading_edge_metabolites).

**Supplementary Table 5.** The results from statistically comparing phenotypic information between the individuals in each cluster versus all other individuals in the VDAART study, as described in the main text. For each variable and each cluster, we included the type of statistical test performed, its associated statistic and p-value, and the type of comparison. For the vitamin D levels and asthma variables we report the mean value and number of True, respectively.

