## Supplementary Table 1 for "The metabolic role of vitamin D in children’s neurodevelopment: a network study"

| Children’s Baseline Characteristics |  | “On Schedule” | “Needs Monitoring” | “Requires further evaluation” |
| --- | --- | --- | --- | --- |
| ASQ-comm |  | 344 (90%) | 23 (6%) | 14 (4%) |
| RACE | White | 115 (33%) | 6 (26%) | 5 (36%) |
|  | Black | 164 (48%) | 13 (57%) | 8 (57%) |
|  | Other | 65 (19%) | 4 (17%) | 1 (7%) |
| SEX | Female | 161 (47%) | 9 0,39130435 | 3 (21%) |
|  | Male | 183 (53%) | 14 0,60869565 | 11 (79%) |
| MATERNAL  EDUCATION | Some college | 78 (23%) | 5 (22%) | 6 (43%) |
|  | High school, Technical school | 91 (26%) | 10 (43%) | 4 (29%) |
|  | College graduate or Graduate school | 131 (38%) | 5 (22%) | 1 (7%) |
|  | Less than high school | 44 (13%) | 3 (13%) | 3 (21%) |
| MATERNAL  MARITAL  STATUS | Separated or divorced | 11 (3%) | 0 (0%) | 0 (0%) |
|  | Married | 169 (49%) | 9 (39%) | 3 (21%) |
|  | Not married and not living together | 85 (25%) | 8 (35%) | 6 (43%) |
|  | Not married but living together | 79 (23%) | 6 (26%) | 5 (36%) |
| SITE | San Diego | 125 (36%) | 6 (26%) | 0 |
|  | Boston | 64 (19%) | 5 (22%) | 9 (64%) |
|  | St Louis | 155 (45%) | 12 (52%) | 5 (36%) |
| TREATMENT | Vitamin D | 176 (51%) | 12 (52%) | 7 (5%) |
|  | Placebo | 168 (49%) | 11 (48%) | 7 (5%) |
| ASTHMA | Asthma True | 86 (25%) | 5 (22%) | 2 (14%) |
|  | Asthma False | 258 (75%) | 18 (78%) | 12 (86%) |
| WHEEZE | Wheeze True | 145 (42%) | 8 (35%) | 8 (57%) |
|  | Wheeze False | 199 (58%) | 15 (65%) | 6 (43%) |
| MATERNAL  ASTHMA | Maternal Asthma True | 142 (41%) | 8 (35%) | 2 (14%) |
|  | Maternal Asthma False | 202 (59%) | 15 (65%) | 12 (86%) |

| **Data Summary** | **Children** | **Mothers** |
| --- | --- | --- |
| **Vitamin D Levels** | Coord Blood +  Year 3 | 32-38 gestation weeks +  1 year after delivery |
| **Metabolomic profiling** | Year 3 | - |
| **ASQ-comm score** | Year 3 | - |
